# Disrupted neural synchrony mediates the relationship between white matter integrity and cognitive performance in older adults

**DOI:** 10.1101/2019.12.29.890228

**Authors:** T. Hinault, M. Kraut, A. Bakker, A. Dagher, S.M. Courtney

**Affiliations:** U1077 INSERM-EPHE-UNICAEN, Caen, FRANCE; Department of Psychological and Brain Sciences, Johns Hopkins University, Baltimore, MD, 21218, USA; Department of Radiology and Radiological Science, Johns Hopkins University School of Medicine, Baltimore, MS, 21287, USA; Department of Psychiatry and Behavioral Sciences, Johns Hopkins University School of Medicine, Baltimore, MD, 21287, USA; F.M. Kirby Research Center, Kennedy Krieger Institute, MD 21205, USA; McConnell Brain Imaging Centre, Montreal Neurological Institute, McGill University, Montréal QC, H3A 2B4, CANADA.; Department of Neuroscience, Johns Hopkins University, MD 21205, USA

**Keywords:** Cognitive control, Aging, EEG, DTI, Connectivity

## Abstract

Our main goal was to determine the influence of white matter integrity on the dynamic coupling between brain regions and the individual variability of cognitive performance in older adults. EEG was recorded while participants performed a task specifically designed to engage working memory and inhibitory processes, and the associations among functional activity, structural integrity, and cognitive performance were assessed. We found that the association between white matter microstructural integrity and cognitive functioning with aging is mediated by time-varying alpha and gamma phase-locking value (PLV). Specifically, older individuals with better preservation of the inferior fronto-occipital fasciculus showed greater task-related modulations of alpha and gamma long-range PLV between the inferior frontal gyrus and occipital lobe, lower local phase-amplitude coupling in occipital lobes, and better cognitive control performance. Our results help delineate the role of individual variability of white matter microstructure in dynamic synchrony and cognitive performance during normal aging, and show that even small reductions in white matter integrity can lead to altered communications between brain regions, which in turn can result in reduced efficiency of cognitive functioning.

**Significance statement:** Cognitive aging is associated with large individual differences, as some individuals maintain cognitive performance similar to that of young adults while others are significantly impaired. We hypothesized that individual differences in white matter integrity would influence the functional synchrony between frontal and posterior brain regions, and cognitive performance in older adults. We found that the association between reduced tract integrity and worse cognitive performance in older adults was mediated by task-related modulations of coupling synchrony in the alpha and gamma bands. Results offer a mechanistic explanation for the neural basis of the variability of cognitive performance in older adults who do not have any clinically diagnosable neuropathology, and for the association between structural network integrity and cognition in older adults.

## 1. Introduction

With aging, an overall decline in cognitive functioning has been observed, with memory and cognitive control being among the first domains to show reduced performance and the greatest changes during aging^1^. Cognitive control is strongly associated with quality of life and autonomy in the elderly^2^, and is usually tested with tasks that involve the active maintenance or updating of goal-relevant information in working memory (WM), and the suppression of competing goal-irrelevant information (inhibition^3,4,5^). Despite an overall decline with age, cognitive function varies widely amongst older individuals. Indeed, while some maintain cognitive performance similar to that of young adults, others are significantly impaired. So far, questions remain about the neural basis of this variability. Investigating the association between white matter microstructure and electroencephalography (EEG) synchrony using connectivity analyses, we aimed to contribute to the understanding of neural mechanisms underlying individual differences in cognitive performance in older adults.

Connectivity between brain regions has been shown to underlie even small differences in cognitive performance between individuals and age groups^6^. Connectivity analyses can be performed at the structural level, by investigating the integrity of white matter tracts, or at the functional level, by studying the temporal attributes of coupling between anatomically separated brain regions. At the structural level, previous studies have found an association between cognitive control performance and white matter tracts connecting frontal and parietal areas in young adults^7^, and age-related reduction of white matter microstructural integrity has been associated with cognitive decline^8^. At the functional level, studies using functional magnetic resonance imaging (fMRI) revealed associations between age–related reductions in fronto-parietal connectivity and reduced effectiveness of cognitive control processes^9^. These results led to the formulation of a disconnection model of cognitive aging^10, 11^, associating interruption of brain connectivity with age-related decline in cognitive performance.

Few studies have investigated how aging effects in structural connectivity relate to functional couplings, and the relationship between structural-functional coupling and cognitive performance has not been systematically investigated^12,13,14,15^. Recently, we used fMRI and diffusion tensor imaging (DTI) to investigate how frontoposterior structural-functional couplings contribute to a better understanding of the inter-individual variability of cognitive control in older adults^16^. Results showed that microstructural integrity of the inferior frontal-occipital fasciculus (IFO) was correlated with task-related frontoposterior effective connectivity between the inferior frontal gyrus (IFG) and occipital lobe, and with the preservation of cognitive control in older adults. However, together with the majority of previous work, this study relied on functional MRI, a method with limited temporal resolution. The influence of age-related structural integrity differences on the temporal dynamics of activity in the engaged networks is still poorly understood, and is critical for identifying the functional neural mechanisms and for guiding strategies for prevention and treatment of age-related cognitive decline.

Oscillatory activity and the task-related modulation of its synchronization are necessary for rapid and selective communication among distant brain regions^17^. Frontoposterior synchrony in the alpha band (8-13 Hz) has been implicated in the effectiveness of inhibitory control^18^, while synchronization in the gamma band (25-100 Hz) has been associated with encoding and information maintenance^19^. Investigation of interactions among these frequency bands revealed modulations of gamma amplitude as a function of the alpha phase within the frontoposterior network. This alpha-gamma phase-amplitude coupling was reduced during encoding and mental manipulation tasks relative to resting-states^20^. These results were interpreted as suggesting that the alpha band regulates information processing through pulsed inhibition (i.e., alternating inhibition and excitation as a function of alpha phase^18, 21^), and that a release of such inhibition is necessary in tasks involving controlled information processing. Aging could be associated with a reduced ability to control and maintain synchronized activity in task-relevant couplings^22^.

Resting-state EEG connectivity partly depends on the underlying white matter tract architecture^23,24,25,26^. With aging, even small changes in white matter integrity might lead to delayed and/or altered communications between brain regions, which could in turn alter cognitive functioning. Although this was suggested by previous work on interhemispheric coherence^27^, the relationship with cognitive performance in older adults has not been investigated. Age-related differences in microstructure could affect the quality and/or the temporal dynamics of communications between brain regions, and contribute to cognitive changes in aging. Results may thus contribute to a mechanistic explanation for why white matter degradation that appears minimal on standard clinical MRI is sometimes associated with substantial changes in cognition with aging^28^.

Here, we investigated the time course of source-reconstructed EEG connectivity in younger and older adults during a cognitive control task, and its association with white matter microstructure and cognitive performance. Participants performed an arithmetic verification task specifically designed to target both WM and inhibition. In line with previous work, we expected a greater arithmetic interference effect in older adults^29^ and a reduction of the facilitated inhibition following WM updating that is observed in young adults^16^, suggesting that these processes are less effective with aging. We also hypothesized that individual differences in microstructural properties of frontoposterior tracts would influence the time-varying functional synchrony of the task-relevant frontoposterior couplings and behavioral interference resolution. We found that the degree to which an older individual’s IFO tract microstructure differs from the microstructural profile of young adults predicts that individual’s task-related synchrony, in both the alpha and gamma bands, between the IFG and the occipital lobe, which in turn predicts facilitated inhibitory control performance. Furthermore, greater synchrony was associated with a reduction of local alpha/gamma phase-amplitude coupling in sensory sites, suggesting that greater long-range synchrony combined with lower local coupling is necessary for inhibition of irrelevant information.

## 2. Methods

### 2.1. Participants

We analyzed data from 40 young adults (18-35) and 40 older adults (60-85; see participant characteristics in Table 1). All participants were right-handed and reported normal or corrected-to-normal vision. Age groups were matched on years of education, reported genders, and arithmetic fluency. Standard MRI exclusion criteria included claustrophobia and the presence of metal in the body. All older adults performed within normal range (i.e., score > 26) on the Montreal Cognitive Assessment (MoCA^30^). No participant had a history of neurological or cognitive disorders, traumatic brain injury, or major psychiatric disorders. Participants were not on any neurologically/psychiatrically active medication at the time of testing (with the exception of a stable anti-depressant regimen for 12 weeks or more). The study was approved by the Johns Hopkins School of Medicine Institutional Review Board. Participants were paid $60. Forty-one older adults were recruited, but one participant was excluded from the analyses due to accuracy below chance in several experimental blocks. Forty-one young adults were recruited, but one participant was excluded from the analyses for completing only one experimental session. Recruitment of young adults continued until the sample size matched the older adult group.

**Table 1:**
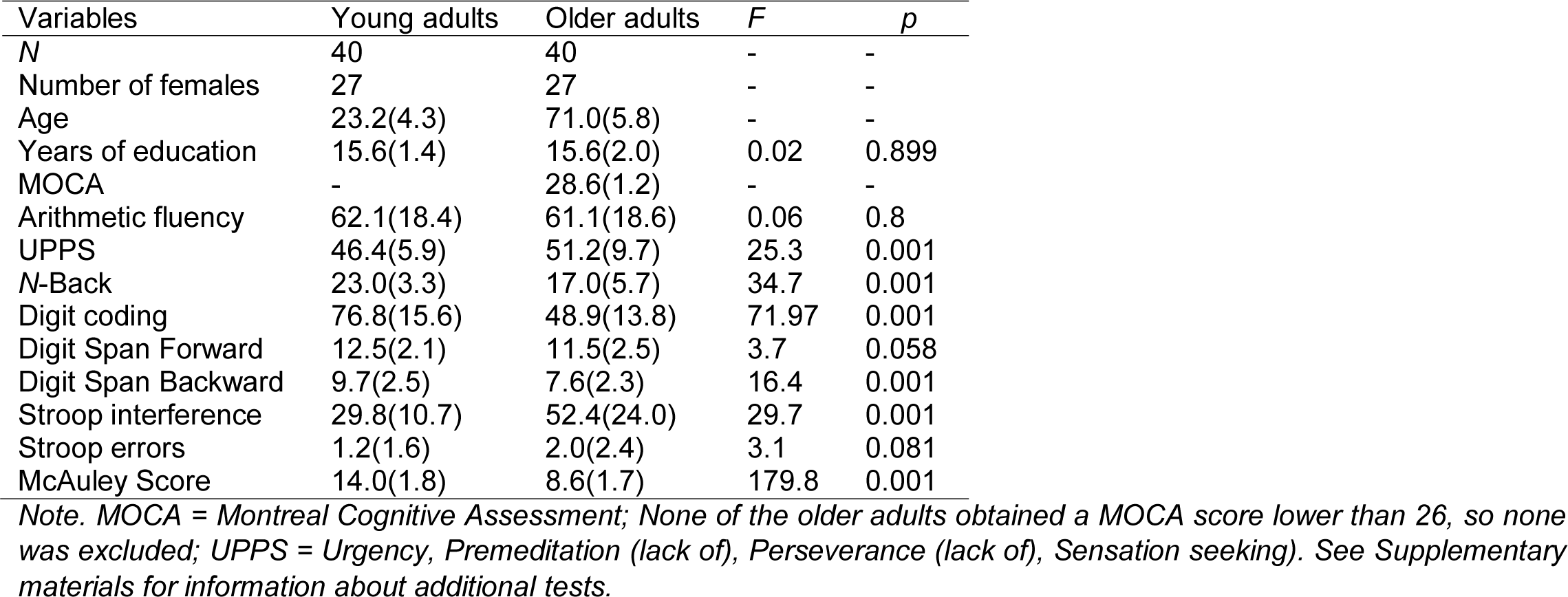
Participants’ characteristics

### 2.2. Experimental Paradigm

The experimental paradigm was identical to our previous study^16^. An arithmetic verification task was implemented in OpenSesame software^31^. Addition or multiplication problems consisted of equations presented in “*a b c*” form, where “*a”* and “*b”* are the operands and “*c”* is the sum or product of the operands. Following previous work^32,33,34^, problems represented all combinations of digits for the “*a”* and “*b”* terms from 2 to 9. One and zero were not included because these problems are solved differently from other problems^35^. The equations 2+2 and 2×2 were not included because the sum and products of these operands are identical, but other equations in which the two operands are the same number (e.g., 3×3) were included.

The operator to be used was cued at the beginning of each trial and was not displayed with the numbers. There was a total of 252 trials divided into 4 blocks of 63 trials (see^16^ for a list of the stimuli used in the task). Half the presented problems (126 trials) were multiplications and half were additions. Half the presented problems had the correct solution to the equation (according to the currently relevant operation) whereas, within the other half, 63 problems were False-related problems (i.e, the proposed solution is correct for the operation that is not currently relevant; e.g., “9 6 15” when the current operation is multiplication) and 63 were False-unrelated problems (i.e., the proposed solution is not correct for either addition or multiplication; e.g., “9 6 14”). False-unrelated problems were created by adding or subtracting 1 to or from the False-related solution. This was done to equalize the distance between the products of False-related and False-unrelated problems, as such distance is known to influence arithmetic processing^36^. This distance was kept constant across operations, and the parity (i.e., even or odd) of operands and results were also equally distributed across the conditions. In each problem, the solution was never one of the two operands (e.g., “2 1 2” was never presented). Also, problems in each condition were equated in numerical size, such that the average sum of the two operands in each condition was the same (i.e., average sum: 11.16). Addition and multiplication equations were presented intermixed to increase the arithmetic interference^37, 38^. Half the presented problems were maintenance problems, in which the cued arithmetic operation did not change within the trial (indicated by a “HOLD” cue between the operator cue and the problem presentation), and half were updating problems, in which the cued arithmetic operation had to be updated to the other operator (“FLIP” cue, indicating a switch from addition to multiplication or vice versa) before the presentation of the operands and proposed solution. The order of the problems was pseudo-randomized to ensure that there was an equal number of repetition and updating of operations (i.e., Hold, Flip), problem type (i.e., addition, multiplication), and condition (i.e., True and False problems) throughout the session, and the block order was counterbalanced across participants. Additional analyses did not reveal significant interactions involving the Operation type factor (*F*s<3.0).

Each trial began with a fixation dot presented for 500ms in the center of the screen (Figure 1A). The operation cue (i.e. a “+” sign for addition problems and an “X” sign for multiplication problems) was then displayed on the computer screen for 1500ms. Following this cue period, a fixation dot was presented with a pseudorandom jitter between 1000ms and 1400ms. Then, the WM instruction (“HOLD”, or “FLIP”) was presented for 1500ms. The “HOLD” cue instructed participants to maintain the cued operation. The “FLIP” cue instructed a change to the other, non-cued operation. Following the WM instruction, a fixation dot was presented with another pseudorandom jitter between 1000ms and 1400ms. Both operands and a proposed answer were then displayed simultaneously on the screen. The proposed answer appeared below the operands. Participants indicated via a gamepad whether the proposed answer was the correct result of the equation given the currently relevant operation, or if the equation was incorrect. Both operands and the proposed answer remained on the screen until participant’s response, or for a maximum duration of 4000ms. Following the problem verification period, a 500ms blank screen was displayed before the next trial started. All participants completed a short training session of 20 problems, different from the experimental stimuli. Instructions emphasized both accuracy and speed. Participants were reminded to keep their eyes fixed upon the white fixation cross and to try to remain as still as possible. There was a pause of approximately 30 seconds after each block of 63 trials. All participants underwent a brief neuropsychological assessment (see supporting information) outside the EEG room.

**Figure 1.**
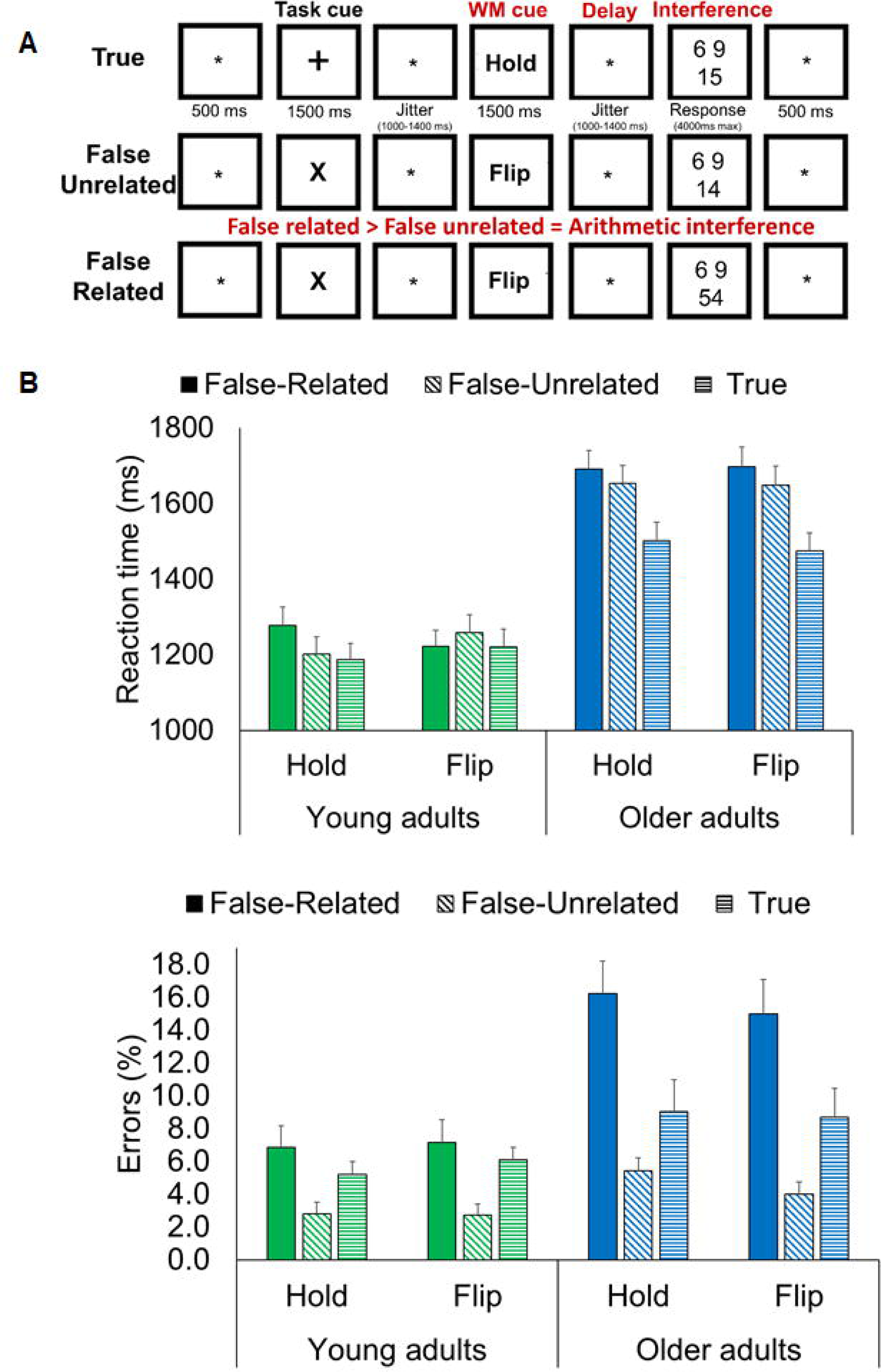
Experimental procedure and behavioral results. A, sequence of events within a trial. B, Mean solution times and percentages of errors for False-related problems, False-unrelated problems, and True problems, as a function of whether a Flip or Hold cue was displayed, in young and older adults. Errors bars represent the standard error of the mean (S.E.M). *p<0.5, **p<0.01, ***p<0.001.

### 2.3. Behavioral analysis

The first trial of each block, and trials with no response or an erroneous response were not included in the analyses. Mean reaction times and percentages of errors were analyzed using 2 (Age: young adults, older adults) × 2 (WM Cue: Hold, Flip) × 3 (Problem type: True, False-related, False-unrelated) mixed-design ANOVAs. To correct for multiple comparisons in ANOVA analyses, Šidák correction^39^ was applied, and Šidák-adjusted p-values are reported. Behavioral interference was defined as the difference in RT for correctly answered False-related – False-unrelated trials.

### 2.4. MRI acquisition

Individuals were scanned with a 3T Phillips Achieva MRI scanner with a 32-channel head coil. Total scanning time was 25 minutes. Following localizer scans, T1-weighted images were acquired with a 3D MP-RAGE sequence: Field of view: 212 × 212 × 172 mm, sagittal orientation, 1 × 1 × 1 mm voxel size, repetition time: 3000ms, echo time: 2ms, flip angle: 8°. Diffusion whole brain images were acquired with the following parameters: 70 axial slices, slice thickness = 2.2 mm, voxel size = 2.21 × 2.21 × 2.2 mm, TR = 2000ms, TE = 71ms, FOV = 212 × 212 × 154 mm. Diffusion gradients were applied along 64 noncollinear directions (b = 1000 s/mm²). Although not presented here, resting-state fMRI data were also collected.

### 2.5. Electroencephalography recording and analyses

EEG activity was recorded inside a Faraday cage with 128 electrodes covering the whole scalp (ActiCHamp, Brain Products, Munich, Germany). Such high-density recording, and adherence to recommended guidelines for source reconstruction and connectivity^40^, were selected to improve source localization accuracy and reduce source leakage^41^. As a projector located outside the EEG room was used to display the task stimuli, we used a photodiode to detect the exact onset time of the stimulus, and this timing information was used to correct for projector-induced delays. Electrode impedance was maintained below 15 kΩ. EEG data were recorded at a sampling rate of 1000Hz. The Fpz electrode was used as ground and the Cz electrodes were used as reference. Artifact and channel rejection (on continuous data), filtering (0.3-100 Hz bandpass, on unepoched data), re-referencing (i.e., using the algebraic average of the left and right mastoid electrodes), time segmentation into epochs, averaging, and source estimation were performed using Brainstorm^42^. In addition, physiological artefacts (e.g., blinks, saccades) were identified and removed through signal-space projection. Power-spectrum density was estimated over the whole recording period to identify bad electrodes and then interpolate data from those electrodes from other neighboring electrodes.

The CapTrack camera system was used to record the spatial positions of the electrodes on the cap. These positions were co-registered to that individual’s anatomy using fiducial points and were used to improve the source reconstruction accuracy. Age-related brain atrophy can introduce changes in the amplitude of event-related activations^43^. Therefore, FreeSurfer^44^ was used to generate cortical surfaces and automatically segment cortical structures from each participant’s T1-weighted anatomical MRI, to account for individual brain atrophy levels during source reconstruction. The EEG forward model was obtained from a symmetric boundary element method (BEM model; OpenMEEG^45, 46^, fitted to the spatial positions of every electrode^47^. A cortically constrained, sLORETA procedure was applied to estimate the cortical origin of scalp EEG signals, and was weighted by a sample estimate of sensor noise covariance matrix^48^ obtained from the baseline periods (i.e., fixation period before the display of the WM cue), in each of the participants, and used to improve data modelling. This method was selected because of reduced localization error and false positive connectivity relative to other source localization methods^49, 50^. The estimated sources were then smoothed (i.e., full width at half maximum: 3mm) and projected in a standard space (i.e., ICBM152 template) for comparisons between groups and individuals, while controlling for differences in native anatomy. This procedure was applied to three time periods of interest: 1) following the WM cue (i.e., from -500ms to 1500ms after the onset of the WM cue), 2) during the delay period (i.e., from 1500 to 3000ms after the onset of the WM cue), and 3) following the problem display (i.e., from - 500ms to 1500ms after the onset of the three numbers). See the supporting information for analyses of the delay period.

### 2.6. Time resolved Phase-locking value (PLV)

Phase-locking analyses^51^ were used to determine the functional coupling between regions of interest PLV estimates the variability of phase differences between two regions across trials. If the phase difference varies little across trials, PLV is close to 1 (i.e., high synchrony between regions) while, with large variability in the phase difference, PLV is close to zero. The range of each frequency band was based on the individual alpha-peak frequency (IAF) observed at posterior sites (i.e., bilateral parietal, parieto-occipital, and occipital sites). Following previous work^52^, the following frequency bands were then considered: Alpha (IAF-2/IAF+2), Gamma1 (IAF+15/IAF+30), and Gamma2 (IAF+31/IAF+80). See supporting information for other frequency bands and additional time-frequency analyses. To preserve timing information while reducing the dimensionality of the data, PLV was estimated at the subject level, across trials. To reduce the dimensionality of the data, the first mode of the principal component analysis (PCA) decomposition of the activation time course in each region of interest (ROI) from the Desikan atlas brain parcellation^53^ was used. The first component, rather than mean activity, was selected to reduce signal leakage^54^. The averages of 100ms sliding time windows were then extracted across the epochs of interest. Data were first analyzed through permutation tests (N=1,000^55^, false discovery rate (FDR) corrected). In the ROIs that showed significant effects in both young and older adults (highlighted in Tables 2 and 3), Age × WM Cue/Problem type (depending on the task time period) × Coupling × Time mixed-design ANOVAs were conducted to test for differences between Flip and Hold trials, between False-related and False-unrelated trials, and aging effects therein. This approach was used to identify interactions between factors and differences between young and older adults in activation time course. Participants’ age and mean grey matter volume were included as covariates in the analyses.

### 2.7 DTI analyses

Preprocessing of diffusion data was done using ExploreDTI^56^ and included the following steps (see also our previous work^16^): (a) Images were corrected for eddy current distortion and participant motion; (b) a non-linear least square method was applied for diffusion tensor estimation, and (c) whole brain DTI deterministic tractography was estimated using the following parameters, for each participant: uniform 2 mm resolution, fractional anisotropy (FA) threshold of 0.2 (limit: 1), angle threshold of 45°, and fiber length range of 50 – 500mm. In line with previous work^57, 58^, the following white matter tracts were delineated in native space: left and right IFO, left and right superior longitudinal fasciculus (SLF), and left and right cingulum bundle (CB). The FA values of each voxel along each tract were extracted for each participant, and fiber bundles were then resampled and normalized to match the length of the average tract length (across group). Following previous work^59^, an FA profile for each tract and each participant was calculated as the average FA of all the voxels within each slice, positioned anteriorly to posteriorly along the tract. A normal young FA profile for each tract was calculated by averaging the FA profiles of the young adults at each point along the tract. Each older adult’s FA was compared to the younger control mean profile. In young adults, normal inter-individual variations in the FA profile were examined by comparing each participant to the FA profile averaged across the profiles of all the other young participants. To quantify the degree to which an individual participant’s tract microstructure was different from the young normal microstructure, we performed a least-squares linear regression for each individual’s profile against the young adults’ profile. This analysis yielded a value for the fit of each participant’s curve for each tract, which reflects how well the shape of the individual’s tract profile matches (i.e., *R*^2^ close to 1) or deviates (i.e., *R^2^* close to 0) from the expected FA profile for that tract. This method was used as a more sensitive measure of the microstructural integrity of the entire tract than average FA value. As an example, a lesion resulting in a localized dip in FA in a segment of a tract could result in an overall normal average FA if FA in other parts of the tract were relatively high, but such a dip would result in a poor profile-fitting^59^.

### 2.8. Correlation, regression and mediation analyses

Correlation analyses were performed between DTI microstructural integrity, PLV synchrony, and behavioral interference to identify task-relevant functional-structural couplings and to specify mediation relationships. Correlations were FDR corrected for multiple comparisons. To limit the number of correlations between EEG and DTI measures, only plausible, long anatomical-distance correlations within hemisphere (i.e., frontoposterior couplings) were assessed. When the same PLV, DTI and behavioral interference measures were correlated within both age groups, the involved variables were selected for mediation analyses to assess the respective contribution and the directionality of the relationships^60^. The tested mediation model (Figure 4) aimed to determine whether PLV mediated the relationships between DTI microstructural integrity and behavioral interference. The analyses estimated the direct effect (Path C) of microstructural integrity on behavior after controlling for PLV synchrony and the indirect effect (Path AB). Direct and indirect effects were computed with bias-corrected bootstrap 95% confidence intervals based on 5,000 bootstrap samples^61^. The mediation was significant if the confidence intervals in Path AB did not cross through zero. Partial mediation was denoted if Path C’ was significant and full mediation was denoted if Path C’ was no longer significant. All analyses were performed using IBM SPSS Statistics for Windows, Version 24.0 with the PROCESS macro for mediation analyses^61^.

### 2.9. Time resolved Phase-amplitude coupling

The coupling between the phase of the alpha frequency and the amplitude of gamma1 and gamma2 was estimated in each condition and time period to assess the relationship between local phase-amplitude coupling and long-range synchrony between brain regions. The time windows of interest were filtered with additional “buffer” windows of 200 ms before and after that window with a bandpass filter for the low and the high frequencies (defined based on individual IAF as described above). The Hilbert transformation was used to extract the instantaneous phase of the slow frequencies and the amplitude of the fast frequencies. To eliminate edge artifacts caused by filtering and applying the Hilbert transformation, the additional buffer windows were removed. Filtering, transforming, and extracting each trial separately, was done to prevent spurious phase-amplitude coupling due to sharp power transitions in the data^19, 62^. To assess time-varying changes of phase-amplitude coupling across the time windows of interest, a 250ms sliding window was used (see supporting information). Correlations were FDR corrected for multiple comparisons.

## 3. Results

### 3.1. Behavioral results

Older adults were slower to respond to the problem verification than young adults, *F*(1,78)=35.25*, p*<.001, *MS*e=225917.16, *np*²=.31 (Figure 1B), and showed larger group variability, *F*(1,78)=4.79*, p*<.032, *MS*e=1111.09, *np*²=.06. The main effect of Problem type (True, False-related, False-unrelated) revealed an arithmetic interference effect, *F*(2,156)=45.04*, p*<.001, *MS*e=8803.31, *np*²=.37. Participants were slower to solve False-related problems than either False-unrelated problems (*F*(1,78)=10.33*, p*=.002, *MS*e=2027.53, *np*²=.12), or True problems (*F*(1,78)=65.98*, p*<.001, *MS*e=32499.96, *np*²=.46). Participants also took more time to solve False-unrelated problems than True problems (*F*(1,78)=37.80*, p*<.001, *MS*e=18292.39, *np*²=.33). The Age x Problem type interaction revealed a smaller difference between young and older adults on True problems compared to False-unrelated problems or False-related, *F*(2,156)=19.61*, p*<.001, *MS*e=3832.77, *np*²=.20. A similar main effect of Problem type was observed with percentages of errors, *F*(2,156)=33.44*, p*<.001, *MS*e=29.54, *np*²=.30. Greater error rates were observed for False-related problems than False-unrelated problems (*F*(1,78)=69.57*, p*<.001, *MS*e=117.94, *np*²=.47), or True problems (*F*(1,78)=17.01*, p*<.001, *MS*e=33.86, *np*²=.18), and False-unrelated problems yielded smaller error rates than True problems, *F*(1,78)=15.75*, p*<.001, *MS*e=12.71, *np*²=.17. Moreover, the Age x Problem type interaction was significant and showed greater differences between age groups for False-related problems relative to False-unrelated or True problems, *F*(2,156)=7.23*, p*=.001, *MS*e=6.39, *np*²=.09. In summary, results are consistent with greater interference effects in older adults on average.

Results replicated the facilitated inhibition following WM updating, and aging effects therein, as observed in our earlier study^16^. Across both age groups, the WM Cue x Problem type interaction revealed that the difference between False-related and False-unrelated problems was significant for Hold trials (*F*(1,78)=5.61, *p*=.020, *MS*e=11.37, *np*²=.07), but not significant for Flip trials (*F*<3.0). Importantly, however, a significant Age x WM Cue x Problem type interaction was found (*F*(2,156)=10.42*, p*<.001, *MS*e=653.58, *np*²=.12). In older adults, inhibition of arithmetic interference was not facilitated following WM updating (Flip trials). Contrasts revealed that the difference between False-related and False-unrelated problems was significant for both Hold and Flip trials in older adults (38ms and 48ms, *F*(2,38)=24.72*, p*<.001, *MS*e=18.76, *np*²=.39, and *F*(2,38)=56.68*, p*<.001, *MS*e=18.33, *np*²=.60, respectively), while in young adults the difference was significant for Hold trials (77ms, *F*(2,38)=10.55*, p*<.001, *MS*e=17.18, *np*²=.22) but not for Flip trials (*F*<3.0). To control for an alternative interpretation of the results in terms of general aging or processing speed, participants’ ages and the digit symbol substitution task scores (see supporting information) were each used as a covariate in ANOVAs (i.e., one ANOVA per covariate, and an additional ANOVA combining these factors). No significant interaction involving age or processing speed was observed (*F*s<2.0). Data were also log-transformed to control for general slowing in baseline performance among older adults, but similar results were observed. No WM Cue x Problem type or Age x WM Cue x Problem type interaction was observed with error rates (*F*s<.5).

To summarize, while young adults showed a reduced arithmetic interference after the cued operation was updated (Flip trials) compared to when the operation was maintained (Hold trials), older adults showed a significant arithmetic interference regardless of the WM cue. These results suggest that in young adults updating from the previously cued operation to the alternative operation (i.e., from addition to multiplication or vice versa) facilitates the inhibition of the arithmetic facts whose retrieval is prompted by the numbers presented in the false-related trials. This facilitation of inhibition, however, is less effective in older adults.

### 3.2. Time-resolved Phase Locking Value (PLV)

Permutation tests were used to identify couplings showing a double dissociation across time between young and older adults (see supporting information for additional analyses). These couplings were included in ANOVA analyses. During the WM cue period (see Table 2, Figure 2), Age x WM cue x Time interactions were observed in the alpha (F(13,1014)=3.05, p<.001, MSe=4.04, np²=.04), and gamma2 bands (*F*(13,1014)=2.99*, p*<.001, *MS*e=3.94, *np*²=.04). These interactions were observed in the right IFG-occipital lobe coupling only (Figure 2A, 2B). No main effects of Age or Cue were observed (*F*s<2.5). Results revealed delayed task-related modulations in older adults relative to young adults. Following the WM cue, in the left IFG-occipital lobe coupling, alpha PLV is transiently greater for Hold than Flip while gamma2 PLV is greater for Flip than Hold. This task-related gamma modulation of PLV emerged later for older adults than young adults, and such difference correlated with behavioral interference in older adults (*r*=-.402, *p*=.010, Figure 2C). During problem verification (see Table 3, Figure 2D), following the Hold cue, an Age x WM cue x Time interaction was observed in the alpha band (*F*(13,1014)=1.91, *p*<.025, *MS*e=8.28, *np*²=.02), for the left IFG-occipital lobe coupling. Again, a delay in task-related modulation of PLV was observed in the older adults. Following the onset of the problem verification phase, alpha PLV was transiently greater for False-related problems than False-unrelated problems in the left IFG-occipital lobe coupling, suggesting alpha synchrony between these brain regions is necessary to inhibit the irrelevant information (*r*=-.467, *p*=.005). This difference in PLV emerged later for older than younger adults. Following the Flip cue, no significant interaction was observed (*F*s<1.5).

**Figure 2.**
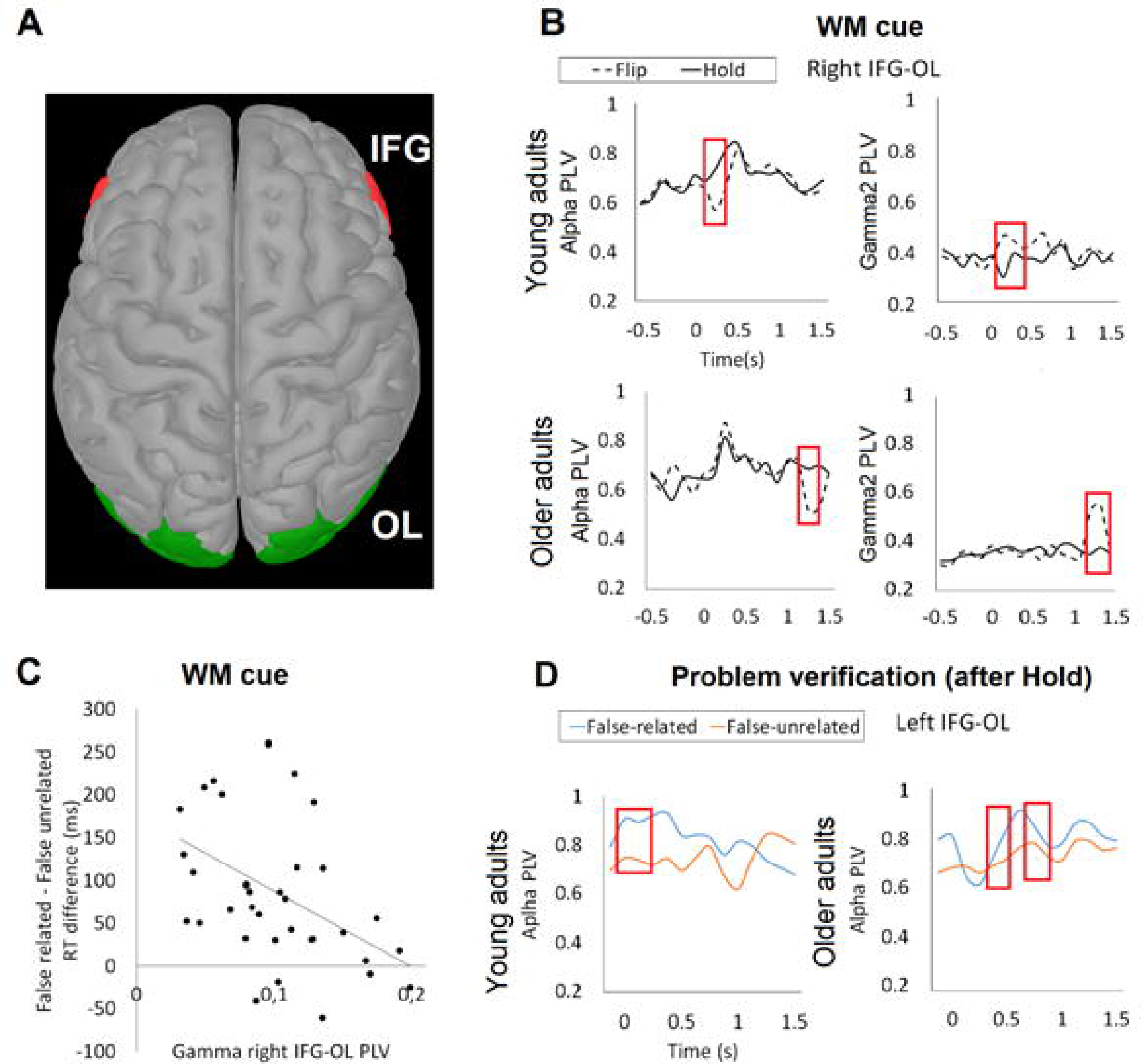
Age-related differences in PLV synchrony in the alpha, theta, and gamma bands. A, Functional coupling was observed between sources in IFG (red) and OL (green). B, Differences in PLV between Flip and Hold trials were observed during the cue period. C, Differences in IFG-OL PLV between Flip and Hold trials was negatively correlated with behavioral interference. D, During the problem verification period, differences in PLV between False-related and False-unrelated trials were observed for trials following Hold cues. Red boxes indicate the time periods with significant contrasts.

**Table 2:**
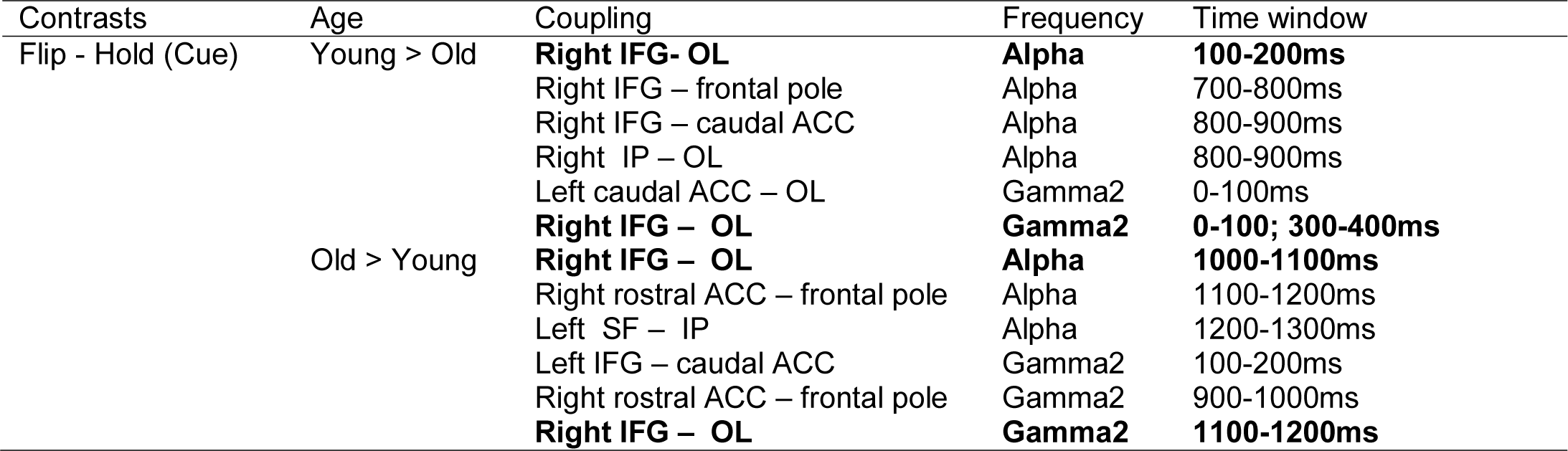
Significant Phase Locking Value couplings following permutation tests for the Flip *minus* Hold difference in the young vs. older adults contrast, as a function of frequency and time following cue onset. ACC= anterior cingulate cortex, IFG=inferior frontal gyrus, OL=occipital lobe, IP = inferior parietal, SF= superior frontal. Results showing a double dissociation (selected for ANOVA analyses) are highlighted. Only results for the alpha and gamma bands are shown here. See supporting information for other frequency bands.

**Table 3:**
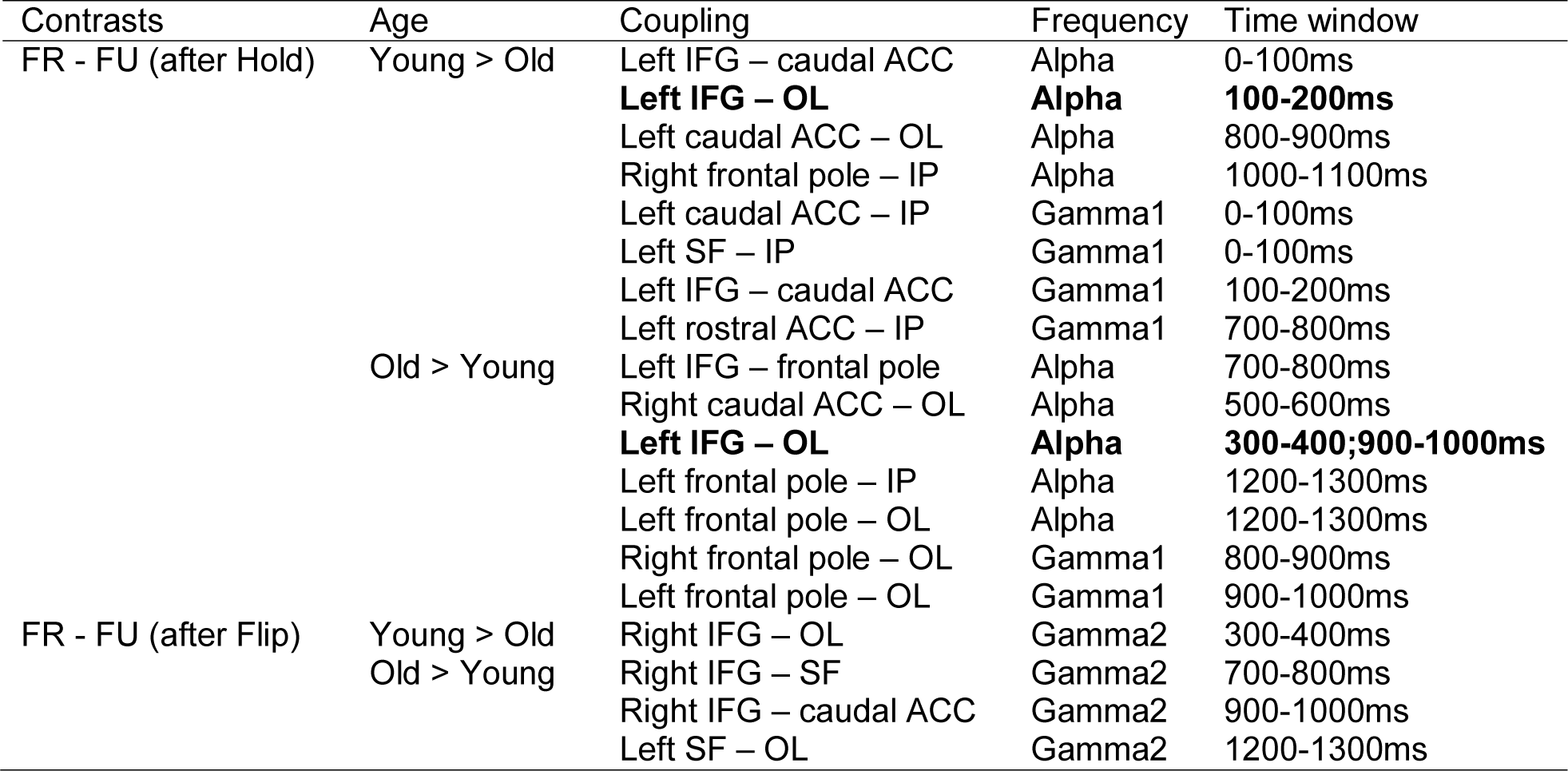
Significant PLV couplings following permutation tests for the Flip *minus* Hold difference and False-related *minus* False-unrelated in the young vs. older adults contrast, as a function of frequency and time following problem onset. ACC= anterior cingulate cortex, IFG=inferior frontal gyrus, OL=occipital lobe, IP = inferior parietal, SF= superior frontal, SP= superior parietal. Results showing a double dissociation (selected for ANOVA analyses) are highlighted. Only results for the alpha and gamma bands are shown here. See supporting information for other frequency bands.

In summary, PLV results revealed task-related changes of functional synchrony during both WM cue and problem verification periods, and age-related differences in both alpha and gamma frequency bands. Converging evidence was found regarding the role of the IFG-OL coupling across time periods and frequency bands. Older adults showed reduced and delayed task-related modulations of synchrony in both frequency bands relative to young adults. Our next goal was to determine the associations among such long-range synchrony, structural connectivity, behavioral performance, and local phase-amplitude coupling.

### 3.3. DTI results

Older adults showed overall smaller FA values than young adults (*F*(1,78)=14.31, *p<*.001, *MS*e=.07, *np*²=.16). This difference in FA values between age groups was similar across tracts (*F*<1.0, Figure 3). In line with behavioral results, variability of tract integrity was overall larger in older adults relative to young adults (*F*(1,78)=4.10, *p<*.046, *MS*e=5.13, *np*²=.05). To look for regional differences in white matter microstructural integrity between young and older adults, additional analyses were conducted with the Segment (frontal first third, middle third, posterior last third) factor on each tract of interest, but no interaction of this factor with age was significant (*F*s<1.0). In young adults, microstructural integrity of the left CB was negatively correlated with the behavioral interference following the Flip cue (*r*=-.463, *p*=.003). In older adults, integrity of the right IFO was negatively correlated with the arithmetic interference following the Flip cue (*r*=-.414, *p*=.008). We then conducted mediation analyses to determine whether the association between tract integrity and behavioral performance in older adults could be mediated by long-range functional synchrony.

**Figure 3:**
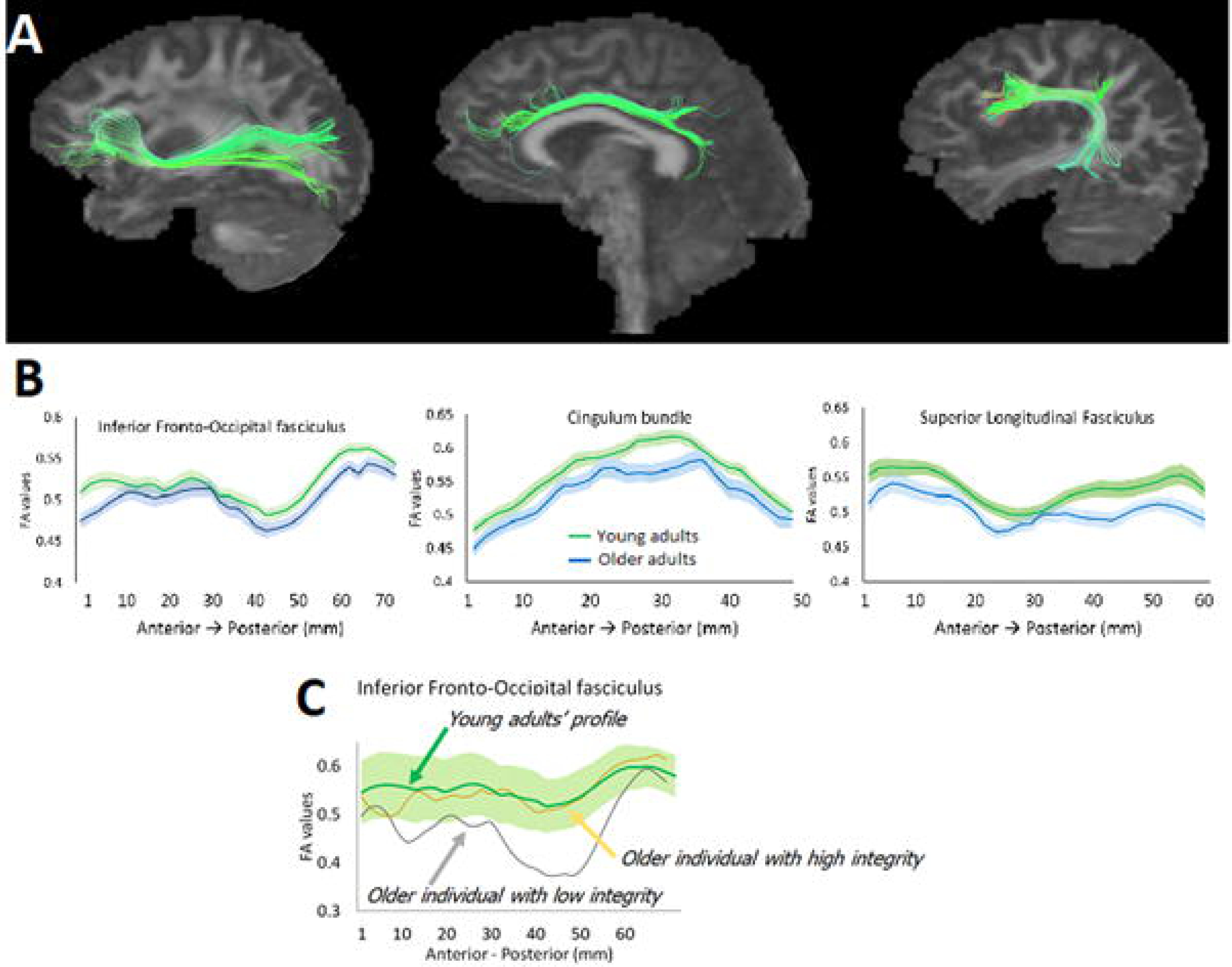
A, The inferior fronto-occipital fasciculus (left), cingulum bundle (middle), and superior longitudinal fasciculus (right). B, White matter microstructural integrity along each tract in young and older adults. C, Example of two older participants with different level of difference from the microstructural profile of young adults in the inferior fronto-occipital tract.

### 3.4. Mediation analyses

As our main goal was to test for potential mediation relationships between microstructural integrity, functional couplings, and behavioral performance, we identified joint correlations (FDR corrected) of PLV measures with behavioral interference and DTI microstructural integrity, and mediation analyses were performed to specify the relationship among these variables. In order to reduce the number of correlations tested, only plausible correlations between structural and functional measures (i.e., frontoposterior variables within the same hemisphere) were investigated. Also, negative correlations between white matter microstructural integrity and PLV synchrony that involved the same structural-functional coupling were not considered because they could reflect indirect (i.e., multiregional) couplings that cannot be directly measured here. Correlation analyses were performed separately for each age group and were FDR corrected for multiple comparisons. Both during the cue and the problem verification periods (Figure 4), mediation analyses revealed that PLV magnitude significantly mediated the relationship between white matter microstructural integrity and behavioral arithmetic interference in older adults. No significant joint relationship was observed in young adults. These analyses involved the microstructural integrity of the IFO tract, alpha and gamma PLV synchrony between IFG and occipital lobe, and arithmetic interference. Effects were observed in the right hemisphere for gamma synchrony following the WM cue, and in the left hemisphere for alpha synchrony during the problem verification. The microstructural integrity of this frontoposterior tract predicted the EEG alpha and gamma synchrony between these brain regions, which in turn predicted behavioral interference. No mediation relationship nor joint correlation was observed in other frequency bands or in homologous regions in the other hemisphere. Age, mean grey matter volume and local IFG and occipital lobe volumes (from the Freesurfer cortical parcellation) were used as covariates in the analyses but no effect was observed on the mediation relationship.

**Figure 4:**
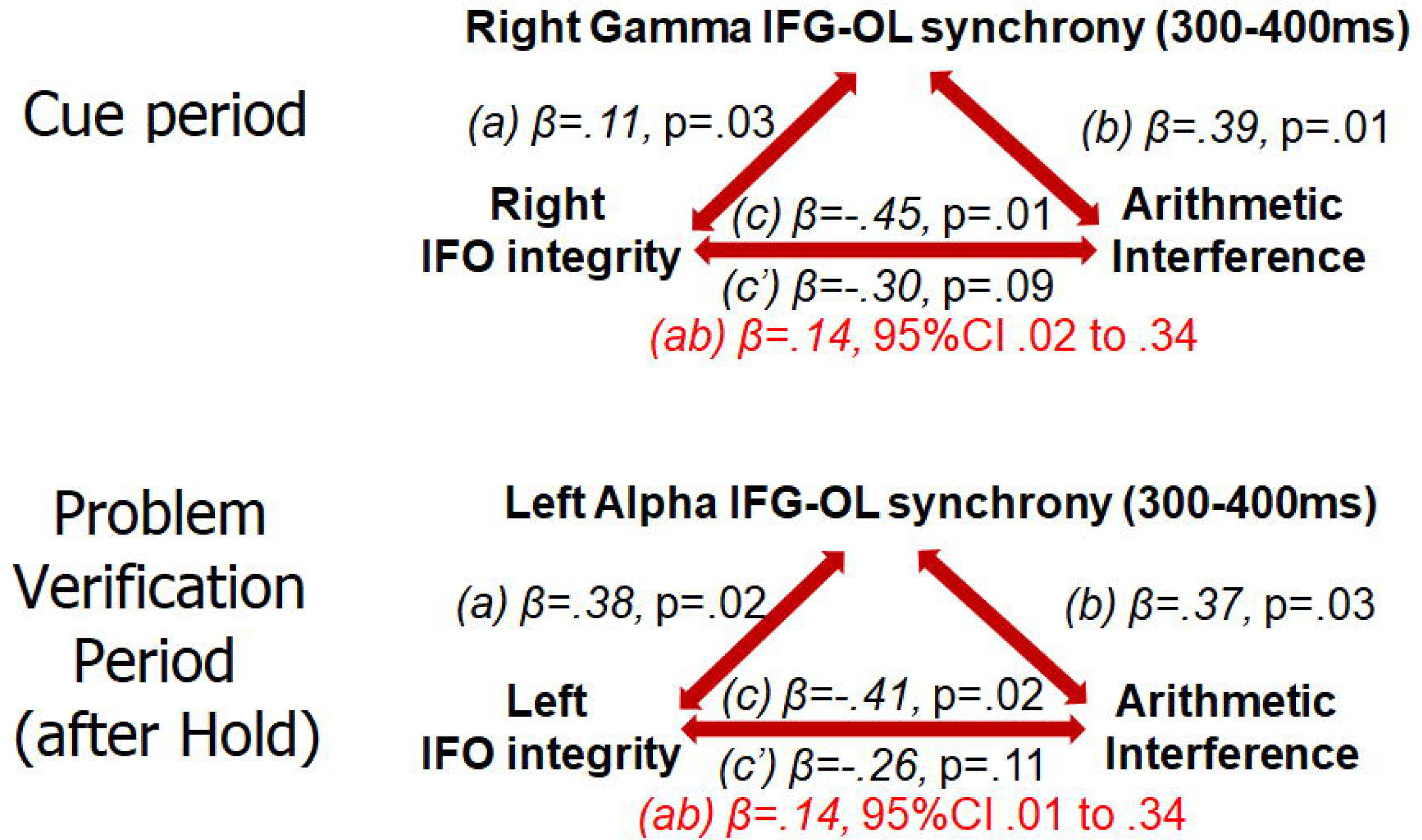
Mediation of white matter integrity effect on arithmetic interference by the functional synchrony between the inferior frontal gyrus (IFG) and the occipital lobe (OL) during the cue period (gamma band) and the problem verification period (alpha band), in older adults.

### 3.5. Time-resolved Phase Amplitude Coupling

Because the relationship between local frequency coupling and long-range synchrony between brain regions may provide insights into the potential role of long-range synchrony in information processing, we also investigated alpha-gamma phase-amplitude coupling in the ROIs showing significant mediation relationship (see supporting information, Figure S1, for additional phase-amplitude coupling analyses). In older adults, correlations were observed following Flip cues between behavioral interference and the alpha-gamma1 phase-amplitude coupling in the right occipital lobe 300-400ms after the WM cue (r=.438, *p*=.005). Such phase-amplitude coupling was also correlated with the right IFG-occipital lobe PLV gamma coupling involved in the mediation analysis (r=-.439, *p*=.005), at the same latency. In other words, lower phase-amplitude coupling between alpha and gamma in OL following the Flip cue was associated with less arithmetic interference (i.e., better cognitive control) and greater functional synchrony between this region and IFG. These results indicate that in False-related trials following a Flip cue, both low local occipital frequency coupling and high IFG-OL long-range synchrony may be needed to update the task-relevant rule and then quickly inhibit irrelevant information.

## Discussion

Our main goal was to determine the influence of white matter integrity on dynamic functional connectivity and cognitive performance during normal aging. To this end, we investigated inhibition following WM updating or maintenance in an arithmetic verification task while EEG activity was recorded, and we quantified individual differences in tract-based DTI measures in young and older adults. First, we replicated the overall decline of facilitated inhibition in older adults relative to younger adults, as older adults showed no reduction of arithmetic interference following updating relative to maintenance cues. EEG activity provided information on the functional synchrony involved (a) when participants were asked to maintain or update the cued arithmetic operation in WM, and (b) when they were using the relevant operation to verify arithmetic equations. We found task-related modulations in frontoposterior couplings to be delayed in older adults relative to young adults. This connectivity and its delayed modulation was specific to the communication between the IFG and the occipital lobe, two areas previously identified as important for inhibitory control in this task. When the association between functional coupling and white matter microstructure was assessed, we found that EEG synchrony mediated the relation between white matter microstructure and the resolution of arithmetic interference. In older adults, relatively preserved white matter microstructural integrity was associated with earlier and greater frontoposterior synchrony and reduced interference effects. These results demonstrate that age-related alterations of white mater microstructure impact long-range dynamic functional synchrony and cognitive performance, and have important implications for our understanding of the variability in cognitive performance in aging.

Investigating oscillatory activity enabled us to determine which frequency bands showed task-related modulations of synchrony, and age-related differences therein. In line with previous work^52^, defining the range of each frequency band based on individual alpha peak frequency provides a sensitive estimate of oscillatory activity and compensates for frequency slowing during aging^63, 64^. Results revealed that the known association between white matter and cognitive performance with aging^10, 65, 66^ is mediated by frontoposterior PLV in the alpha and gamma bands when cognitive control processes are engaged. This mediation was observed during both WM cue and problem verification phases, and involved synchronization between the IFG and occipital lobe regions. IFG activity has been reported when rule updating and maintenance is required^67,68,69,70^, and its coupling with the occipital lobe was previously observed with the same task in fMRI^16^, suggesting a major role for the interplay between rule updating and sensory processing. The present study shows that, in contrast with a disconnection account, there is no absolute interruption of network structure or communication, but rather reduced integrity leads to reduced synchrony between brain regions, and slower transitions for task-related modulations of that synchrony. Such functional network alterations appear to lead in turn to altered cognitive performance in situations where it is necessary to quickly and effectively disengage for no-longer relevant rules or to inhibit processing of irrelevant information. There is not a presence or absence of communication between brain regions in aging, but rather a continuum, with varying degrees of slowed and inefficient communication, corresponding to the continuum observed in cognitive performance.

Results also provide new and interesting findings on the relationship between frontoposterior PLV and local occipital phase-amplitude coupling measures. 300-400ms following the onset of the Flip WM cue, gamma synchrony between IFG and occipital lobe was negatively correlated with the occipital lobe alpha-gamma phase-amplitude coupling. The reduced phase-amplitude coupling was also associated with better behavioral performance as shown by smaller interference effects. These results suggest that cognitive performance following a rule update is normally associated with a transient disruption of local sensory phase-amplitude coupling together with increased long-range phase synchrony in young adults. The reduction of this apparent dissociation during aging is related to cognitive impairment, suggesting impaired WM rule updating and, relatedly, impaired or slowed changes in local sensory processing necessary to inhibit information that has just become irrelevant. A reduction in phase-amplitude coupling in occipital lobe following the WM updating cue could be necessary to quickly reconfigure processing according to communication with IFG through long-range phase synchrony. Such combination of lower local phase-amplitude coupling and higher frontoposterior PLV might facilitate fast and effective disengagement of the no-longer relevant cued operation to allow retrieval of the now-relevant one. In contrast with WM maintenance^19^, where a positive association between theta PLV and theta-gamma phase amplitude coupling was observed, updating might require low local coupling between faster and slower frequencies so that the phase of the faster frequency related to the newly-relevant information can synchronize with that of distant brain regions. An older individual with preservation of such rapid switches of synchrony and local couplings would then be expected to have preservation of the facilitated inhibition following updating that is observed in young adults, as we have observed.

In addition to the previously demonstrated age-related differences of frontoposterior structural and functional couplings^8, 15, 16^, we also found that young and older adults differ in the task-related temporal dynamics of network engagement. Source reconstruction of EEG activity with individual anatomical information, although not as anatomically accurate as fMRI, uniquely combines excellent temporal and good spatial resolutions^71^. PLV analyses revealed similar task-related modulations of alpha and gamma frontoposterior couplings in young and older adults, but effects were observed later in older adults relative to young adults. This result suggests that even if there is no complete interruption of network synchrony, such delayed and potentially noisier communication no longer enables effective network reconfiguration during WM updating, which in turn is associated with greater behavioral arithmetic interference during the subsequent problem verification.

Following the display of the Flip cue, frontoposterior couplings showed a transient decrease of synchrony in the alpha band and a corresponding transient increase of synchrony in the gamma band. This pattern of results is consistent with the previously proposed gating role of the alpha band and suggests a regulation of information processing in a top-down way through pulsed inhibition^18, 21^. The results are also consistent with previous interpretations of long-range synchrony in the gamma band as reflecting information processing and neural communication^72, 73, 75^. Although several studies interpreted oscillatory activity in the gamma band as reflecting mostly local processing^75^, these studies mainly investigated amplitude, rather than phase. Other studies observed long-range gamma synchrony^72^ in association with working memory processing^52, 76^. In the current study, relative to young adults, older adults showed smaller and slower-evolving task-related effects in the gamma band, which is consistent with lower effectiveness of updating processes during task completion.

These findings are constrained by several limitations. First, because of the specific investigation of PLV couplings between frontal and posterior brain regions, indirect communications through additional brain regions could not be detected even though they might be associated with cognitive performance. Although this partly reflects the limitation of the PLV method used, the method also ensures a proper, specific association with identifiable white matter tracts between brain regions. Second, although the methods used were selected, as in multiple magnetoencephalography (MEG) and EEG connectivity analysis papers^54, 77, 78^, to limit the risk of false-positives and to control for signal leakage across ROIs, the anatomical parcellation used may be sub-optimal for detecting interactions between smaller brain regions. However, the relatively low spatial resolution of EEG precludes finer-grained analyses, and this method has been validated on EEG data^79^, giving confidence in the reported results. Finally, the absence of association between cognitive performance and white matter microstructure, and the absence of a significant mediation relationship in young adults likely reflect the significantly lower within-group variability (but see^80^ for results on the association between white matter microstructure and processing speed).

Although cognitive trajectories with aging can strongly vary across individuals^81, 82^, the neural mechanisms underlying these differences are still not well understood. Here, we showed that considering the close relationship between structural and time-varying functional connectivity at different frequencies might be critical to understanding individual differences in cognitive functioning and to be able to detect these even in the absence of clear neuroanatomical lesions or pathological cognitive decline on standardized neuropsychological tests. While this interaction between structural and dynamic connectivity is not considered by previous theoretical frameworks of brain aging^83, 84^, the current study demonstrates its importance. Individuals with a relative preservation of microstructural integrity showed a better functional synchrony between brain regions and similar cognitive performance to that of young adults. Conversely, others with greater structural alterations showed reduced functional couplings, delayed modulation of those couplings, and did not benefit from WM updating when processing arithmetic interference. This is especially true for long-range frontoposterior white matter fibers that are more susceptible to reduced integrity with age^85^. These results point to a possible mechanistic account of the commonly reported finding of decreased “speed of processing” in older adults^86^ and in other populations that have reduced white matter structural integrity such as traumatic brain injury^87^ and multiple sclerosis^88^.

## Acknowledgements

We would like to thank Eda Incekara, Claire Narang, Daniella Needleman, Pranit Singh, Phillip Sumardi, and Travis Kroeker, for their help with data collection and preprocessing. Carrie Speck helped with participant recruitment and screening. Thomas Hinault designed the research, collected and analyzed the data, and wrote the paper. Michael Kraut helped analyze the data and write the paper. Arnold Bakker helped analyze the data and write the paper. Alain Dagher helped analyze the data and write the paper. Susan Courtney helped design the research, analyze the data, and write the paper.

This work was supported by the Albstein Research Foundation, the Fahs-Beck fund to Research and Experimentation, and the William and Ella Owens Medical Research Foundation.

