## Supplementary material for "Disrupted neural synchrony mediates the relationship between white matter integrity and cognitive performance in older adults": Table 1

Table 1: Participants’ characteristics

| Variables | Young adults | Older adults | *F* | *p* |
| --- | --- | --- | --- | --- |
| *N* | 40 | 40 | - | - |
| Number of females | 27 | 27 | - | - |
| Age | 23.2(4.3) | 71.0(5.8) | - | - |
| Years of education | 15.6(1.4) | 15.6(2.0) | 0.02 | 0.899 |
| MOCA | - | 28.6(1.2) | - | - |
| Arithmetic fluency | 62.1(18.4) | 61.1(18.6) | 0.06 | 0.8 |
| UPPS | 46.4(5.9) | 51.2(9.7) | 25.3 | 0.001 |
| *N*-Back | 23.0(3.3) | 17.0(5.7) | 34.7 | 0.001 |
| Digit coding | 76.8(15.6) | 48.9(13.8) | 71.97 | 0.001 |
| Digit Span Forward | 12.5(2.1) | 11.5(2.5) | 3.7 | 0.058 |
| Digit Span Backward | 9.7(2.5) | 7.6(2.3) | 16.4 | 0.001 |
| Stroop interference | 29.8(10.7) | 52.4(24.0) | 29.7 | 0.001 |
| Stroop errors | 1.2(1.6) | 2.0(2.4) | 3.1 | 0.081 |
| McAuley Score | 14.0(1.8) | 8.6(1.7) | 179.8 | 0.001 |

*Note. MOCA = Montreal Cognitive Assessment; None of the older adults obtained a MOCA score lower than 26, so none was excluded; UPPS = Urgency, Premeditation (lack of), Perseverance (lack of), Sensation seeking). See Supplementary materials for information about additional tests.*
