## Supplementary material for "Disrupted neural synchrony mediates the relationship between white matter integrity and cognitive performance in older adults": Table 2

| Contrasts | Age | Coupling | Frequency | Time window (ms) |
| --- | --- | --- | --- | --- |
| Flip - Hold (Cue) | Young > Old | **Right IFG- OL** | **Alpha** | **100-200** |
|  |  | Right IFG – frontal pole | Alpha | 700-800 |
|  |  | Right IFG – caudal ACC | Alpha | 800-900 |
|  |  | Right IP – OL | Alpha | 800-900 |
|  |  | Left caudal ACC – OL | Gamma2 | 0-100 |
|  |  | **Right IFG – OL** | **Gamma2** | **0-100; 300-400** |
|  | Old > Young | **Right IFG – OL** | **Alpha** | **1000-1100** |
|  |  | Right rostral ACC – frontal pole | Alpha | 1100-1200 |
|  |  | Left SF – IP | Alpha | 1200-1300 |
|  |  | Left IFG – caudal ACC | Gamma2 | 100-200 |
|  |  | Right rostral ACC – frontal pole | Gamma2 | 900-1000 |
|  |  | **Right IFG – OL** | **Gamma2** | **1100-1200** |

Table 2: Significant Phase Locking Value couplings following permutation tests for the Flip *minus* Hold difference in the young vs. older adults contrast, as a function of frequency and time following cue onset. ACC= anterior cingulate cortex, IFG=inferior frontal gyrus, OL=occipital lobe, IP = inferior parietal, SF= superior frontal. Results showing a double dissociation (selected for ANOVA analyses) are highlighted. Only results for the alpha and gamma bands are shown here. See supporting information for other frequency bands.
