## Supplementary material for "Disrupted neural synchrony mediates the relationship between white matter integrity and cognitive performance in older adults": Table 3

Table 3: Significant PLV couplings following permutation tests for the Flip *minus* Hold difference and False-related *minus* False-unrelated in the young vs. older adults contrast, as a function of frequency and time following problem onset. ACC= anterior cingulate cortex, IFG=inferior frontal gyrus, OL=occipital lobe, IP = inferior parietal, SF= superior frontal, SP= superior parietal. Results showing a double dissociation (selected for ANOVA analyses) are highlighted. Only results for the alpha and gamma bands are shown here. See supporting information for other frequency bands.

| Contrasts | Age | Coupling | Frequency | Time window (ms) |
| --- | --- | --- | --- | --- |
| FR - FU (after Hold) | Young > Old | Left IFG – caudal ACC | Alpha | 0-100 |
|  |  | **Left IFG – OL** | **Alpha** | **100-200** |
|  |  | Left caudal ACC – OL | Alpha | 800-900 |
|  |  | Right frontal pole – IP | Alpha | 1000-1100 |
|  |  | Left caudal ACC – IP | Gamma1 | 0-100 |
|  |  | Left SF – IP | Gamma1 | 0-100 |
|  |  | Left IFG – caudal ACC | Gamma1 | 100-200 |
|  |  | Left rostral ACC – IP | Gamma1 | 700-800 |
|  | Old > Young | Left IFG – frontal pole | Alpha | 700-800 |
|  |  | Right caudal ACC – OL | Alpha | 500-600 |
|  |  | **Left IFG – OL** | **Alpha** | **300-400;900-1000** |
|  |  | Left frontal pole – IP | Alpha | 1200-1300 |
|  |  | Left frontal pole – OL | Alpha | 1200-1300 |
|  |  | Right frontal pole – OL | Gamma1 | 800-900 |
|  |  | Left frontal pole – OL | Gamma1 | 900-1000 |
| FR - FU (after Flip) | Young > Old | Right IFG – OL | Gamma2 | 300-400 |
|  | Old > Young | Right IFG – SF | Gamma2 | 700-800 |
|  |  | Right IFG – caudal ACC | Gamma2 | 900-1000 |
|  |  | Left SF – OL | Gamma2 | 1200-1300 |
