## Supporting information for "Disrupted neural synchrony mediates the relationship between white matter integrity and cognitive performance in older adults"

**Methods**

*Additional tests:* The Stroop task (Stroop, 1935) was used to measure inhibition. In the first part of this test, words designating colors were written in black, and participants were instructed to read the words as quickly and accurately as possible. In the second part, words designating colors were written in different color inks, and participants were instructed to name the color of the ink as quickly and accurately as possible. An index of interference was created by subtracting the time to complete part A from the time to complete part B for each participant, with greater differences indicating larger interference (e.g., less cognitive control). The number of errors during part B (i.e., reading the word instead of naming the ink color) was also used as a measure of inhibitory control. Participants were also administered the short version of the UPPS scale (Urgency, Premeditation (lack of), Perseverance (lack of), Sensation seeking; Cyders et al., 2014) to measure impulsivity. To assess working memory (WM), forward and backward digit spans were measured (Terman, 1916). In this test, the participant heard a sequence of numerical digits and recalled the sequence in backward or in forward order, with increasingly longer sequences being tested in each trial. The final numbers of items correctly repeated forward and backward (i.e., the testing ended were participants made two consecutive errors on a given sequence length) were used as indicators of an individual’s abilities for, respectively, WM storage and manipulation/updating. The auditory *2*-back test (Kirchner, 1958) was used as an additional measure of WM updating. In this test, participants heard a continuous sequence of spoken letters and indicated whether or not a presented letter was the same as heard two temporal positions earlier. We measured the number of correct responses out of 28 trials. Processing speed was assessed using the WAIS-III digit symbol substitution task (Wechsler, 1997), which consisted of nine digit-symbols pairs followed by a list of digits. Participants wrote as many corresponding symbols under each digit as they could in two minutes. To measure arithmetic abilities, the French Kit test (French et al., 1963) was used. In this test, participants solved as many basic arithmetic problems (e.g., 53-18 = X) as possible in 6 minutes. Finally, the McAuley equation was used to assess cardiorespiratory fitness (McAuley et al., 2012). The equation is based on age, sex, body mass index, resting heart rate, and self-reported physical activity level.

*Time-frequency analyses:* Time-frequency decomposition was performed on source data using a Hilbert transform that measures the envelope of the signal after filtering in every frequency band. The range of each frequency band was based on the individual’s peak alpha frequency observed at posterior sites (i.e., bilateral parietal, parieto-occipital, and occipital sites). Following Toppi et al. (2018), the following frequency bands were then considered: Delta (IAF-8/IAF-6), Theta (IAF-6/IAF-2), Alpha (IAF-2/IAF+2), Beta (IAF+2/IAF+14), Gamma1 (IAF+15/IAF+30), and Gamma2 (IAF+31/IAF+80). The signal was then normalized to compensate for the 1/*f* decrease in power, as this decrease can be greater in older adults relative to young adults (e.g., Voytek et al., 2015), and event-related synchronization/desynchronization was assessed relative to the baseline period. Nonparametric permutation tests (*N*=1,000; Maris & Oostenveld, 2007) were used to statistically test for differences between Flip and Hold trials, together than between False-related and False-unrelated trials, and were FDR corrected for multiple comparisons. To reduce the dimensionality of the data, the first mode of the principal component analysis (PCA) of the activation time course in each Regions-Of-Interest (ROI) from the Desikan atlas brain parcellation (Desikan et al., 2006) was used for the statistics, with a minimum activation duration thresholding of 20 ms. The PCA, rather than the mean, was selected to reduce signal leakage (e.g., Sato et al 2018). In order to reduce the number of comparisons, only couplings within a given hemisphere and involving regions previously observed to be activated in this paradigm (i.e., bilateral caudal anterior cingulate cortex (ACC), frontal pole, inferior parietal, occipital lobe, IFG pars triangularis, superior frontal, and rostral ACC ; Hinault et al., 2019) were included in ANOVA analyses. EEG power and PLV were also analyzed during the delay period (i.e., from 1500ms to 3000ms after the onset of the WM cue).

*Time resolved Phase-amplitude coupling (PAC).* To assess time-varying changes of PAC across the time windows of interest, a 250ms sliding window was used. To statistically test for differences between Flip and Hold trials, between False-related and False-unrelated trials, and aging effects therein, Age x WM Cue/Problem type (depending on the task time period) x Region x Time mixed-design ANOVAs were conducted, and contrasts were FDR corrected for multiple comparisons. Participant’s age and mean grey matter volume were included as covariates in the analyses. Only significant interactions were reported.

**Results**

*EEG source results (power).* We investigated the effect of age on inhibition and WM updating by contrasting the difference between Flip and Hold cue, and between False-related and False-unrelated problems, in young and older adults. During the cue period, results were observed in every frequency band (see Table S1), but mostly in the theta, alpha, and gamma1 bands. Group comparisons revealed greater differences between Flip and Hold cues in young adults than in older adults in the right hemisphere, while older adults showed significant differences in both left and right hemispheres. Moreover, a larger number of effects were observed in the theta and alpha bands were observed in older adults relative to young adults. Results involved the right superior frontal (SF), right superior parietal (SP), right occipital lobe and right IFG in young adults, while activations of bilateral IFG and middle frontal regions were observed in older adults.

During the problem verification phase (see Table S2), following the Hold cue, differences between False-related and False-unrelated problems were mainly observed in the alpha and gamma1 bands in young adults, while differences mainly involved the alpha band in older adults. Moreover, activations were mainly observed in the left hemisphere in young adults and in the right hemisphere for older adults. Effects were greater in the left IFG and middle frontal regions in young adults, while the ACC, SP and inferior parietal (IP) showed larger differences in older adults. Following the Flip cue, differences between False-related and False-unrelated problems were observed in the gamma bands in young adults, while additional activations in the delta band were observed in older adults. Moreover, activations were mainly observed in the right hemisphere in young adults and in both left and right hemispheres for older adults. The right IFG and SP regions showed larger differences in young adults, while bilateral OFC, occipital lobe and caudal middle frontal regions were observed in older adults.

In summary, several brain regions showed significant activity differences between Flip and Hold cues, and between False-related and False-unrelated problems. Between young and older adults, differences were observed in the frequency of activation (i.e., alpha and gamma bands in young adults; lower gamma and higher delta/beta activations in older adults) and lateralization (i.e., less lateralized activity in older adults relative to young adults). Moreover, greater frontal activations were observed in older adults relative to young adults.

*Time-resolved Phase Locking Value* (PLV). Functional couplings between brain regions were investigated in each frequency band. During the WM cue period, Age x WM Cue x Coupling x Time interactions were observed in the delta (*F*(143,11154)=1.46*, p*<.001, *MS*e=2.69, *np*²=.02), theta (*F*(143,11154)=1.25*, p*=.022, *MS*e=1.65, *np*²=.02), alpha (*F*(143,11154)=1.71*, p*<.001, *MS*e=2.32, *np*²=.02), and gamma2 (*F*(143,11154)=1.42, *p*<.001, *MS*e=2.42, *np*²=.02) bands. Significant Age x Time interactions also revealed delayed effects in older adults relative to young adults such that the difference between PLV following Flip vs. Hold cues emerged later for older adults in the delta (*F*(13,1014)=2.20*, p*=.008, *MS*e=.01, *np*²=.027) and gamma2 (*F*(13,1014)=1.97*, p*=.020, *MS*e=1.78, *np*²=.03) bands. Overall (see main document Table 2, Figure 2B), contrasts revealed a larger number of couplings in the delta band and involving frontal regions in older adults relative to young adults, while young adults showed greater activity in the alpha band and more frontoposterior couplings.

During the delay period (Table S3), Age x WM Cue x Coupling x Time interactions were observed in the beta (F(143,11154)=2.18, p=.009, MSe=3.25, np²=.03), and gamma1 (F(143,11154)=2.41, p=.003, MSe=3.55, np²=.03) bands. In the theta band, a WM Cue x Coupling x Time (F(143,11154)=1.31, p=.009, MSe=1.78, np²=.02) interaction was observed but did not involve the age factor. No main effect of Age or Cue were observed in these frequency bands (Fs<3.0). Contrasts revealed a larger number of couplings involving the theta band in older adults relative to young adults. Moreover, relative to young adults, older adults showed fewer frontoposterior couplings and a larger number of frontofrontal couplings.

During problem verification (see main document Table 3, Figure 2C and D), following the Hold cue, Age x Problem type x Coupling x Time interactions were observed in the delta (*F*(143,11154)=1.25*, p*=.025, *MS*e=7.80, *np*²=.02), alpha (*F*(143,11154)=1.79*, p*=.040, *MS*e=9.98, *np*²=.02), beta (*F*(143,11154)=1.25*, p*=.006, *MS*e=2.42, *np*²=.02), and gamma1 (*F*(143,11154)=1.25*, p*<.001, *MS*e=2.30, *np*²=.01) bands. No main effect of age was observed (*F*s<1.0) but the main effect of Problem type was significant in each of these bands (delta, *F*(1,78)=10.52*, p*=.002, *MS*e=.02, *np*²=.12; alpha, *F*(1,78)=22.41*, p*<.001, *MS*e=.01, *np*²=.22; beta, *F*(1,78)=37.43*, p*<.001, *MS*e=.01, *np*²=.32; gamma1, *F*(1,78)=30.22*, p*<.001, *MS*e=.01, *np*²=.28), with higher PLV for False-related problems compared to False-unrelated problems. Relative to young adults, contrast revealed that older adults showed a larger number of significant couplings in the delta band and fewer couplings involving the gamma band, but also showed a lower number of frontoposterior couplings relative to local (i.e., frontal or posterior) couplings.

Following the Flip cue, Age x Problem type x Coupling x Time interactions were observed in the delta (*F*(143,11154)=1.26*, p*=.019, *MS*e=7.32, *np*²=.02), theta (*F*(143,11154)=1.30*, p*=.009, *MS*e=5.82, *np*²=.02), and gamma2 (*F*(143,11154)=1.31*, p*=.008, *MS*e=6.63, *np*²=.02) bands. No main effect of Age was observed in these frequency bands (*F*s<1.0), but a main effect of Problem Type was observed in the theta band (*F*(1,78)=4.97*, p*=.029, *MS*e=.01, *np*²=.06), with greater PLV for False-related relative to False-unrelated problems. Compared to young adults, older adults showed a larger number of significant couplings in the delta and theta bands and fewer couplings involving the gamma band. Moreover, older adults showed a larger number of local couplings relative to frontoposterior couplings. Additional analyses were conducted with age and mean grey matter volume as covariate, but, with the exception of effects in the delta band when age was added as covariate (during the WM cue period), the reported results remained significant. However, a main effect of age (*F*(1,78)=85.98*, p*<.001, *MS*e=.03, *np*²=.52) was observed in grey matter volume, with lower overall grey matter volume in older adults than in young adults.

*Time-resolved Phase Amplitude Coupling (PAC).* Functional interaction between lower and higher frequency bands were investigated by conducting alpha-gamma and theta-gamma PAC analyses for each time period, separately for gamma1 and gamma2 (Table S4; Figure S1). During the cue period, a significant omnibus interaction was observed for the alpha-gamma1 PAC (F(36,2808)=1.57, p=.017, MSe=8.26, np²=.02). Moreover, an Age x Time interaction revealed significantly delayed effects in older adults relative to young adults (F(3,468)=3.39, p=.003, MSe=3.34, np²=.042). Contrasts revealed significantly greater PLV for Flip than Hold in the left frontal pole, and left inferior parietal, in young adults than in older adults, while right rostral ACC showed greater PLV for Hold cues. Moreover, greater differences were observed in left caudal ACC, right occipital lobe, and right inferior parietal in older adults relative to in young adults. During the delay period, a significant omnibus interaction was observed for the theta-gamma2 PAC (F(36,2808)=1.43, p=.048, MSe=1.49, np²=.02). Moreover, a main effect of the cue was observed (F(1,78)=6.01, p=.017, MSe=.01, np²=.07), with lower PAC for Flip compared to Hold. Contrasts revealed significantly greater Flip vs. Hold difference in the left caudal ACC, left frontal pole, right occipital lobe, and right IFG in older adults than in young adults.

During problem verification, following the Hold cue, a significant omnibus interaction was observed for the alpha-gamma1 PAC (*F*(8,624)=2.04*, p*=.040, *MS*e=1.07, *np*²=.03). Moreover, a main effect of Problem type was observed (*F*(1,78)=6.68*, p*=.012, *MS*e=6.92, *np*²=.08), with reduced PAC for False-related compared to False-unrelated problems. Contrasts revealed greater differences between False-related and False-unrelated problems in right IFG, right inferior parietal, and left occipital lobe in older adults relative to young adults. Following the Flip cue, no interaction was observed but a main effect of age was found for theta-gamma2 PAC (*F*(1,78)=4.10*, p*=.046, *MS*e=.01, *np*²=.05), with lower PAC for young adults related to older adults. No joint correlation was observed between PAC, PLV and arithmetic interference. Additional analyses were conducted with age and mean grey matter volume as covariate, but the reported results remained significant.

In summary, significant differences in alpha-gamma and theta-gamma PAC were observed between young and older adults during task completion. Differences in alpha-gamma PAC were observed following the Flip/Hold cue and during problem verification, while effects involved the theta-gamma PAC during the delay period. Significant effects were only observed during the cue period in young adults, while effects were observed in every task period in older adults.

Table S1: Significantly activated ROIs (based on the Desikan atlas) for the Flip *minus* Hold difference in the young vs. older adults contrast during the cue period, as a function of frequency and time following cue onset. ACC=anterior cingulate cortex; IFG=inferior frontal gyrus, IP=inferior parietal, SP=superior parietal, OL=occipital lobe, SF= superior frontal, OFC= orbitofrontal cortex, IT=inferior temporal, MT= middle temporal, ST=superior temporal, MF=middle frontal

| *Contrasts* | Age | Regions | Frequency | Time |
| --- | --- | --- | --- | --- |
| *Flip vs. Hold (Cue)* | Young > Old | Right IP | Delta | 0-60ms |
|  |  | Left paracentral | Delta | 0-114ms |
|  |  | Right SP | Delta | 0-194ms |
|  |  | Right caudal MF | Theta | 114-202; 360-454ms |
|  |  | Right precentral | Theta | 372-482ms |
|  |  | Right SF | Theta | 0-106ms |
|  |  | Right supramarginal | Theta | 288-368ms |
|  |  | Right OL | Alpha | 106-194ms |
|  |  | Right IFG triangularis | Alpha | 162-232ms |
|  |  | Left precentral | Alpha | 0-106ms |
|  |  | Right rostral MF | Alpha | 146-240ms |
|  |  | Left ST | Alpha | 0-170ms |
|  |  | Left supramarginal | Alpha | 1046-1094ms |
|  |  | Right caudal MF | Beta | 106-138ms |
|  |  | Left OL | Beta | 396-482ms |
|  |  | Right MT | Beta | 262-286ms |
|  |  | Left IT | Gamma1 | 726-756ms |
|  |  | Right OL | Gamma1 | 98-146ms |
|  |  | Left MT | Gamma1 | 738-770ms |
|  |  | Right IFG triangularis | Gamma1 | 382-406ms |
|  |  | Left OL | Gamma2 | 522-584ms |
|  |  | Right OFC | Gamma2 | 404-438ms |
|  |  | Right precentral | Gamma2 | 90-114ms |
|  |  | Right SF | Gamma2 | 612-652ms |
|  |  | Right SP | Gamma2 | 398-422ms |
|  | Old > Young | Right postcentral | Delta | 1062-1140ms |
|  |  | Left rostral MF | Delta | 44-312ms |
|  |  | Right rostral MF | Delta | 130-312ms |
|  |  | Right supramarginal | Delta | 660-824ms |
|  |  | Left IT | Theta | 490-600ms |
|  |  | Right insula | Theta | 326-600ms |
|  |  | Left IFG opercularis | Theta | 540-604ms |
|  |  | Right IFG opercularis | Theta | 420-546ms |
|  |  | Left rostral MF | Theta | 256-336ms |
|  |  | Right SP | Theta | 620-674ms |
|  |  | Right ST | Theta | 718-788ms |
|  |  | Left caudal MF | Alpha | 514-616ms |
|  |  | Left IT | Alpha | 318-412ms |
|  |  | Left MT | Alpha | 296-360ms |
|  |  | Left IFG opercularis | Alpha | 548-620ms |
|  |  | Left postcentral | Alpha | 502-548ms |
|  |  | Left rostral MF | Alpha | 754-832ms |
|  |  | Left SF | Alpha | 462-526ms |
|  |  | Right ST | Alpha | 74-280;938-1008ms |
|  |  | Right IP | Beta | 762-794ms |
|  |  | Left IT | Gamma1 | 474-514ms |
|  |  | Right IFG triangularis | Gamma1 | 280-304ms |
|  |  | Right precuneus | Gamma1 | 288-320ms |
|  |  | Left SF | Gamma1 | 304-352;398-430ms |
|  |  | Right OL | Gamma2 | 336-360ms |
|  |  | Left OFC | Gamma2 | 138-162ms |
|  |  | Right SF | Gamma2 | 912-936ms |
|  |  | Right SP | Gamma2 | 738-754;1124-1156ms |

Table S2: Significantly activated ROIs (based on the Desikan atlas) for False-related *minus* False unrelated difference in the young vs, older adults contrasts following Hold and Flip cues, as a function of frequency and time following problem onset. ACC=anterior cingulate cortex; IFG=inferior frontal gyrus, IP=inferior parietal, SP=superior parietal, OL=occipital lobe, SF= superior frontal, OFC= orbitofrontal cortex, IT=inferior temporal, MT= middle temporal, ST=superior temporal, MF=middle frontal, SF= superior frontal.

| Contrasts | Age | Regions | Frequency | Time |
| --- | --- | --- | --- | --- |
| FR vs FU (after Hold) | Young > Old | Left IFG opercularis | Delta | 68-282ms |
|  |  | Right SF | Delta | 182-512ms |
|  |  | Left caudal MF | Theta | 146-374ms |
|  |  | Left rostral MF | Theta | 610-792ms |
|  |  | Right caudal MF | Alpha | 578-692ms |
|  |  | Right MT | Alpha | 0-210ms |
|  |  | Left IFG triangularis | Alpha | 146-326;668-750ms |
|  |  | Right ST | Alpha | 0-228ms |
|  |  | Left caudal MF | Beta | 1126-1192ms |
|  |  | Left rostral MF | Beta | 512-610ms |
|  |  | Left IT | Gamma1 | 424-446ms |
|  |  | Right rostral MF | Gamma1 | 390-472ms |
|  |  | Left SP | Gamma1 | 276-358ms |
|  |  | Right Medial OFC | Gamma2 | 708-758ms |
|  |  | Left IFG triangularis | Gamma2 | 406-465ms |
|  | Old > Young | Right IFG orbitalis | Delta | 446-660ms |
|  |  | Right IP | Theta | 1236-1350ms |
|  |  | Right ST | Theta | 734-1060ms |
|  |  | Right rostral ACC | Alpha | 1070-1170ms |
|  |  | Right IFG orbitalis | Alpha | 376-442ms |
|  |  | Left rostral MF | Alpha | 294-376ms |
|  |  | Right IP | Beta | 708-792ms |
|  |  | Right SP | Beta | 610-726ms |
|  |  | Right IP | Gamma1 | 346-446ms |
| FR vs FU (after Flip) | Young > Old | Right postcentral | Delta | 840-1088ms |
|  |  | Right supramarginal | Theta | 906-972ms |
|  |  | Right paracentral | Alpha | 232-298ms |
|  |  | Right insula | Beta | 358-390ms |
|  |  | Right OL | Beta | 298-348ms |
|  |  | Right IFG opercularis | Gamma1 | 244-308ms |
|  |  | Right precuneus | Gamma1 | 216-264ms |
|  |  | Right SP | Gamma1 | 446-512ms |
|  |  | Left IP | Gamma2 | 446-496ms |
|  |  | Left medial OFC | Gamma2 | 80-112ms |
|  |  | Left rostral ACC | Gamma2 | 308-358ms |
|  |  | Right ST | Gamma2 | 626-660ms |
|  | Old > Young | Right OL | Delta | 758-890ms |
|  |  | Left lateral OFC | Delta | 0-512ms |
|  |  | Right lateral OFC | Delta | 0-504ms |
|  |  | Left MT | Delta | 644-758ms |
|  |  | Left postcentral | Delta | 430-660ms |
|  |  | Right IT | Theta | 610-692ms |
|  |  | Left SP | Theta | 130-228ms |
|  |  | Left medial OFC | Alpha | 162-374;718-782ms |
|  |  | Left SP | Alpha | 374-472ms |
|  |  | Left IT | Beta | 308-374ms |
|  |  | Left medial OFC | Beta | 914-978ms |
|  |  | Left ST | Beta | 216-264ms |
|  |  | Left fusiform | Gamma1 | 626-692ms |
|  |  | Right SP | Gamma1 | 430-504ms |
|  |  | Right caudal MF | Gamma2 | 1094-1142ms |
|  |  | Right OL | Gamma2 | 578-626ms |
|  |  | Right IFG triangularis | Gamma2 | 330-380ms |
|  |  | Left SP | Gamma2 | 244-292ms |

Table S3: Significant Phase Locking Value couplings for the Flip minus Hold difference in the young vs. older adults contrast, as a function of frequency and time following cue offset. ACC= anterior cingulate cortex, IFG=inferior frontal gyrus, OL=occipital lobe, IP = inferior parietal, SF= superior frontal.

| Contrasts | Age | Coupling | Frequency | Time window |
| --- | --- | --- | --- | --- |
| Flip – Hold (Cue) | Young > Old | Right rostral ACC – frontal pole | Delta | 100-200ms |
|  |  | Right IFG– frontal pole | Theta | 100-200ms |
|  |  | Left caudal ACC – OL | Theta | 400-500ms |
|  |  | Right IP - OL | Theta | 800-900ms |
|  | Old > Young | Left caudal ACC – IP | Delta | 0-100ms |
|  |  | Left IFG – caudal ACC | Delta | 0-100ms |
|  |  | Left caudal ACC – IP | Theta | 0-100ms |
| Flip – Hold (delay) | Young > Old | Right IFG – frontal pole | Theta | 1500-1600ms |
|  |  | Right frontal pole – OL | Theta | 1900-2000ms |
|  |  | Right IFG – OL | Theta | 1900-2000ms |
|  |  | Left IFG – IP | Theta | 2400-2500ms |
|  |  | Right IFG – IP | Beta | 1600-1700ms |
|  |  | Left rostral ACC – IP | Gamma1 | 2200-2300ms |
|  | Old > Young | Left rostral ACC – IP | Theta | 1500-1600ms |
|  |  | Right caudal ACC – SF | Theta | 1900-2000ms |
|  |  | Left IFG – caudal ACC | Theta | 1900-2000; 2100-2200ms |
|  |  | Right SF – caudal ACC | Theta | 2100-2200ms |
|  |  | Right frontal pole – caudal ACC | Theta | 2400-2500ms |
|  |  | Left IFG – Frontal pole | Beta | 2300-2400ms |
|  |  | Left IFG – Frontal pole | Gamma1 | 2300-2400ms |
| FR - FU (after Hold) | Young > Old | Right caudal ACC – SF | Delta | 300-400; 400-500ms |
|  | Old > Young | Left IFG – caudal ACC | Delta | 200-300ms |
|  |  | Left IFG – frontal pole | Delta | 200-300ms |
|  |  | Right caudal ACC – OL | Delta | 700-800ms |
|  |  | Left rostral ACC – IP | Delta | 700-800ms |
|  |  | Left caudal ACC – SF | Delta | 700-800ms |
|  |  | Left SF – IP | Delta | 700-800ms |
|  |  | Right SF – OL | Delta | 700-800ms |
|  |  | Left IP – OL | Delta | 1000-1100ms |
|  |  | Left rostral ACC – SP | Delta | 1300-1400ms |
|  |  | Right IFG – SF | Delta | 1300-1400ms |
| FR - FU (after Flip) | Young > Old | Right IP – OL | Delta | 300-400ms |
|  |  | Right caudal ACC – SF | Theta | 700-800ms |
|  | Old > Young | Right IFG – IP | Delta | 200-300ms |
|  |  | Right IFG – caudal ACC | Delta | 400-500ms |
|  |  | Left IP – OL | Delta | 500-600ms |
|  |  | Right IP – OL | Delta | 800-900ms |
|  |  | Right SF – OL | Theta | 600-700ms |
|  |  | Right IP – OL | Theta | 800-900ms |
|  |  | Left caudal ACC – OL | Theta | 900-1000ms |
|  |  | Left SF – IP | Theta | 1000-1100ms |

Table S4: Significant phase-amplitude coupling differences between Flip and Hold cues for young adults and older adults. ACC= anterior cingulate cortex, IFG=inferior frontal gyrus, OL=occipital lobe, IP = inferior parietal.

| Contrasts / Coupling | Age | Region | Time window |
| --- | --- | --- | --- |
| Flip - Hold (Cue) | Young > Old | Left frontal pole | 250-500ms |
| Alpha-gamma1 |  | Left IP | 250-500ms |
|  |  | Right rostral ACC | 1250-1500ms |
|  | Old > Young | Right OL | 0-250ms |
|  |  | Left caudal ACC | 250-500ms |
|  |  | Right IP | 0-250ms |
| Flip – Hold (delay) | Old > Young | Left caudal ACC | 1500-1750ms |
| Theta-gamma2 |  | Left frontal pole | 1750-2000ms |
|  |  | Right OL | 1500-1750ms |
|  |  | Right IFG | 2500-2750ms |
| FR - FU (after Hold) | Old > Young | Right IP | 500-750ms |
| Alpha-gamma1 |  | Left OL | 500-750ms |
|  |  | Right IFG | 250-500ms |

Figure S1: Age-related difference in phase-amplitude coupling between alpha, theta, and gamma bands in occipital sites. Differences were observed during the cue, delay and problem verification periods.

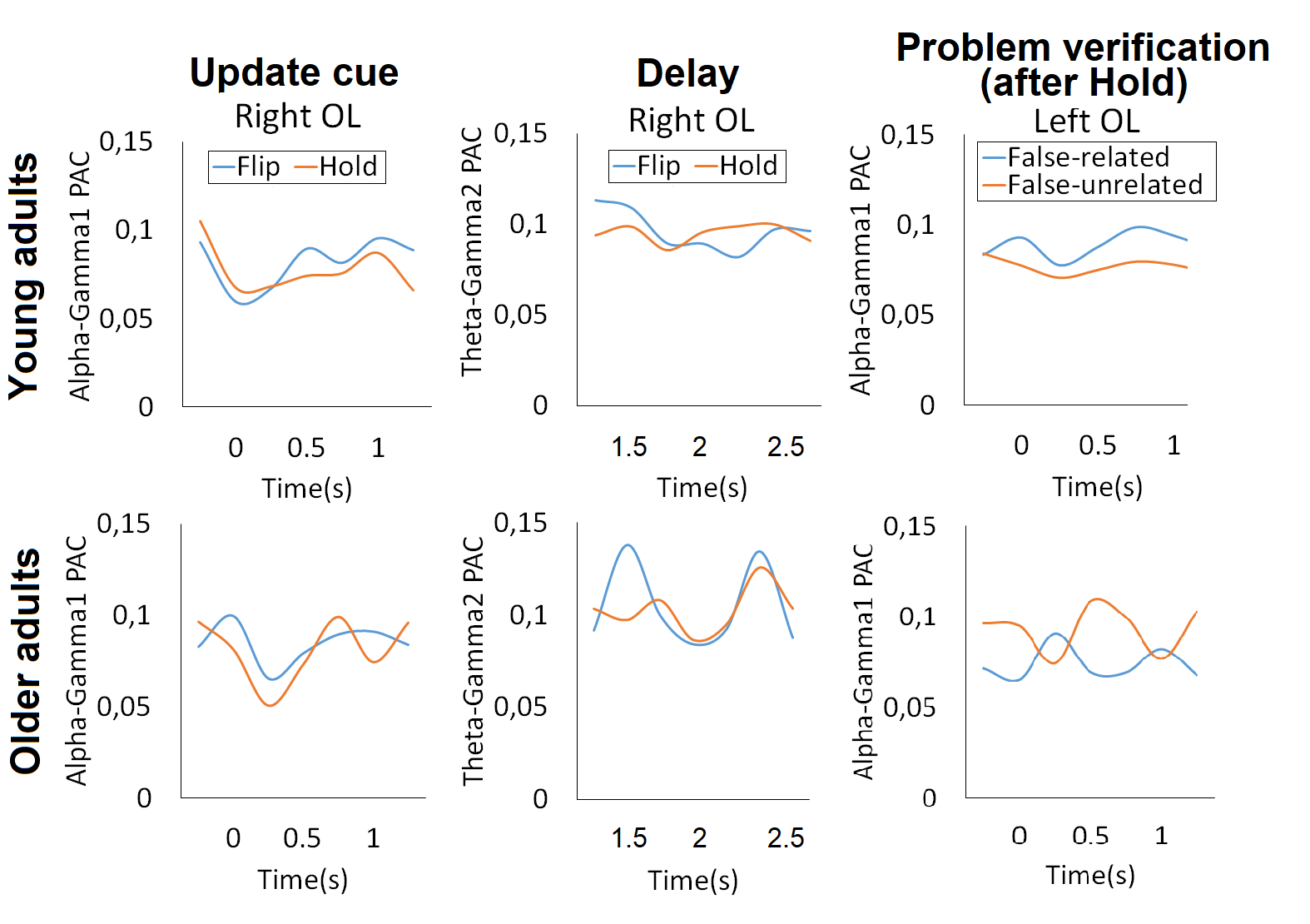
